# The early development and physiology of *Xenopus laevis* tadpole lateral line system

**DOI:** 10.1101/2020.04.21.052969

**Authors:** Valentina Saccomanno, Heather Love, Amy Sylvester, Wen-Chang Li

**Author notes:** Correspondence, School of Psychology and Neuroscience, University of St Andrews, UK.

## Abstract

*Xenopus laevis* has a lateral line mechanosensory system throughout its full life cycle. Previous studies of the tadpole lateral line system revealed that it may play a role in escape behaviour. In this study, we used DASPEI staining to reveal the location of tadpole lateral line neuromasts. Destroying these neuromasts with neomycin resulted in loss of escape responses in tadpoles. We then studied the physiology of anterior lateral line in immobilised tadpoles. Activating the neuromasts behind one eye could evoke asymmetrical motor nerve discharges when the tadpole was resting, suggestive of turning/escape, followed by fictive swimming. When the tadpole was already producing fictive swimming however, anterior lateral line activation reliably led to the termination of swimming. The anterior lateral line had spontaneous afferent discharges at rest, and when activated showed typical adaptation. There were also efferent activities during tadpole swimming, the activity of which was loosely in phase with ipsilateral motor nerve discharges, implying modulation by the motor circuit from the same side. Calcium imaging experiments located sensory interneurons in the primary anterior lateral line nucleus in the hindbrain. Future studies are needed to reveal how sensory information is processed by the central circuit to determine tadpole motor behaviour.

**Summary statement:** Activating tadpole anterior lateral line evokes escape responses followed by swimming and halts ongoing swimming. The afferent and efferent activities and sensory interneuron locations in the hindbrain are reported.

## Introduction

Aquatic vertebrates including fish and Anura use the distributed mechanosensory lateral line (LL) system to sense hydrodynamic disturbances (Bleckmann H and Zelick R, 2009; Coombs S and Zaidi ZH, 2012; Kroese AB et al., 1978; Montgomery JC et al., 1997; Stewart WJ et al., 2013). The LL system provides an important sensory mode for animals to conduct rheotaxis (facing into and swimming against a current) (Akanyeti O et al., 2016; Haehnel-Taguchi M et al., 2018; Montgomery JC, Baker CF and Carton AG, 1997; Oteiza P et al., 2017; Simmons AM et al., 2004), predator avoidance and escape (McHenry MJ et al., 2009; Roberts A et al., 2009; Stewart WJ, Cardenas GS and McHenry MJ, 2013) and prey detection (Claas B and Munz H, 1996; Nagiel A et al., 2008). There are three types of LL receptor: mechanosensory neuromasts, pit organs (Northcutt RG, 1992) and electroreceptive ampullary organs. Superficial neuromasts, embedded in the epidermis, are found in both amphibians and fish, whereas canal neuromasts, protected in grooves, are exclusive to fish (Bleckmann H and Zelick R, 2009; Northcutt RG, 1992; Quinzio S and Fabrezi M, 2014). Hair cells in the centre of neuromasts act as mechanotransducers (Northcutt RG, 1992) while kinocilia embedded inside the cupula bend with water current flow, allowing the detection of water current directions (Nagiel A, Andor-Ardo D and Hudspeth AJ, 2008; Strelioff D and Honrubia V, 1978).

The larval Zebrafish LL system has seven nerves: the supraorbital, infraorbital, mandibular, middle, opercular, otic, middle, and occipital LL nerves (Raible DW and Kruse GJ, 2000). Several studies have been carried out in zebrafish larvae to test the role of the LL system behaviourally. The LL system has been shown to be critical in sensing predation (McHenry MJ, Feitl KE, Strother JA and Van Trump WJ, 2009; Stewart WJ, Cardenas GS and McHenry MJ, 2013), predator evasion (Olszewski J et al., 2012) and rheotaxis (Oteiza P, Odstrcil I, Lauder G, Portugues R and Engert F, 2017; Suli A et al., 2012). LL physiology has been studied by directly recording hair cells (Corey DP and Hudspeth AJ, 1983; Lv C et al., 2016; Olt J et al., 2016; Ricci AJ et al., 2013) and primary afferent neurons (Glowatzki E and Fuchs PA, 2002; Keen EC and Hudspeth AJ, 2006; Liao JC, 2010; Liao JC and Haehnel M, 2012).

Arrangement of the LL system is largely conserved in amphibians (Schlosser G, 2002). All anuran tadpoles have two orbital and three mandibular LL nerves, as well as trunk and tail LL nerves (Quinzio S and Fabrezi M, 2014). The orbital and trunk LL nerves are largely conserved between anuran tadpole species, whilst the mandibular LL nerve exhibits more interspecies variation (Quinzio S and Fabrezi M, 2014). Fish possess an LL system throughout their lifecycle, whereas presence and retention of the LL system in amphibians is dependent on the extent to which they are aquatic. The size of neuromast, and number and organisation of hair-cells, varies between species of Anura (Quinzio S and Fabrezi M, 2014). In *Xenopus*, neuromasts first appear in stage 32 tadpoles (Nieuwkoop PD and Faber J, 1956; Roberts A, Feetham B, Pajak M and Teare T, 2009). Both receptor organs and sensory neurons of the LL develop from progenitor cells deposited by placodes (Schlosser G, 2002; Schlosser G and Northcutt RG, 2000). The five LL placodes of *Xenopus;* anterodorsal, anteroventral, middle, supratemporal, and posterior; derive from the dorsolateral placode (Schlosser G and Northcutt RG, 2000). Each LL placode generates a single LL nerve, as well as at least one sensory ridge or migrating primordia, which differentiates into neuromasts (Schlosser G, 2002). The ganglion cells are produced first, followed by elongated sensory ridges, and finally neuromasts (Schlosser G and Northcutt RG, 2000). *Xenopus* also retains the LL system through metamorphosis into adulthood (Shelton PM, 1970; Strelioff D and Honrubia V, 1978). The adult *Xenopus* has ~200 neuromasts in lines over its head and body, organised into ‘stitches’ of 3-12 neuromasts. Each neuromast comprises 30-60 ciliated hair-cells covered by a gelatinous cupula (Strelioff D and Honrubia V, 1978). In early tadpole stages however, it is thought that the cupula has not yet formed (Roberts A, Feetham B, Pajak M and Teare T, 2009). Activation of the lateral line system in tadpoles can initiate escape responses followed by swimming (Roberts A, Feetham B, Pajak M and Teare T, 2009) and rheotactic behaviour in older tadpoles (Haehnel-Taguchi M, Akanyeti O and Liao JC, 2018; Simmons AM, Costa LM and Gerstein HB, 2004).

The spinal cord of *Xenopus laevis* tadpoles at stage 37/38 is one of the simplest and arguably best understood among all vertebrates (Roberts A et al., 2012; Roberts A et al., 2010). Tadpole sensory modalities are fairly limited at stage 37/38. Only mechanosensory systems and the light-sensing pineal eyes are functional. The LL system of the hatchling *Xenopus* tadpole provides a very simple sensory system for water current detection (Roberts A, Feetham B, Pajak M and Teare T, 2009). Other senses including ocular vision, hearing, taste and pain are not developed. The simplicity of tadpole sensory and motor systems presents the animal as an excellent model for studying sensory information integration and motor decision-making (Koutsikou S et al., 2018; Roberts A et al., 2019). In this study, we investigated the neurophysiology of the tadpole LL system.

## Materials and Methods

Mating between pairs of adult *Xenopus laevis* was induced regularly by injections of human chorionic gonadotropin (HCG, 1000 U/ml, Sigma, UK) into the dorsal lymph sacs. Procedures for HCG injections comply with UK Home Office regulations. All experiment procedures on tadpoles were approved by the Animal Welfare Ethics Committee (AWEC) of the University of St Andrews.

To visualise the LL neuromasts in tadpole skin, 2-[4-(dimethylamino)styryl]-N-ethylpyridinium iodide (DASPEI), was obtained from Sigma-Aldrich UK. Small batches of stock were made up in distilled H_2_0 prior to experiments and stored at 4°C in darkness to prevent photobleaching. An assay by Pisano, et al. (Pisano GC et al., 2014) was adapted for use in the present study. A solution of 0.4 mg/ml DASPEI was made in distilled water. Before staining, the stock solution was diluted with equal volume of saline (concentrations in mM: NaCl, 115; KCl, 3; CaCl_2_, 2; NaHCO_3_, 2.4; MgCl_2_, 1; HEPES, 10; pH adjusted to 7.4 with NaOH). Tadpoles were placed in the final 0.2 mg/ml solution for 30 minutes in darkness on a rocking bed, then placed in a saline bath for 5 minutes to wash off excess dye. To identify and count stained neuromasts, a stereomicroscope was modified with the addition of a barrier filter and monochromatic LED light source at 450nm to allow fluorescent imaging. Tadpoles were placed in saline between two recessed slides. Both sides of the head and trunk were observed and photographed using an eyepiece camera and DinoCapture imaging software (Dino-Lite Europe). To enhance visualisation of the neuromasts, the original fluorescent microscopy images were converted to grey scale images by enhancing the red and yellow colour channels (Fig. 1 A). All images were edited using FIJI for ImageJ or Corel PHOTO PAINT, and neuromast measurements were also taken with FIJI (Schindelin J et al., 2012). All experiments were performed on newly hatched prefeeding *Xenopus laevis* larvae between developmental stages 32-42, defined by Nieuwkoop and Faber (Nieuwkoop PD and Faber J, 1956).

**Fig. 1.**
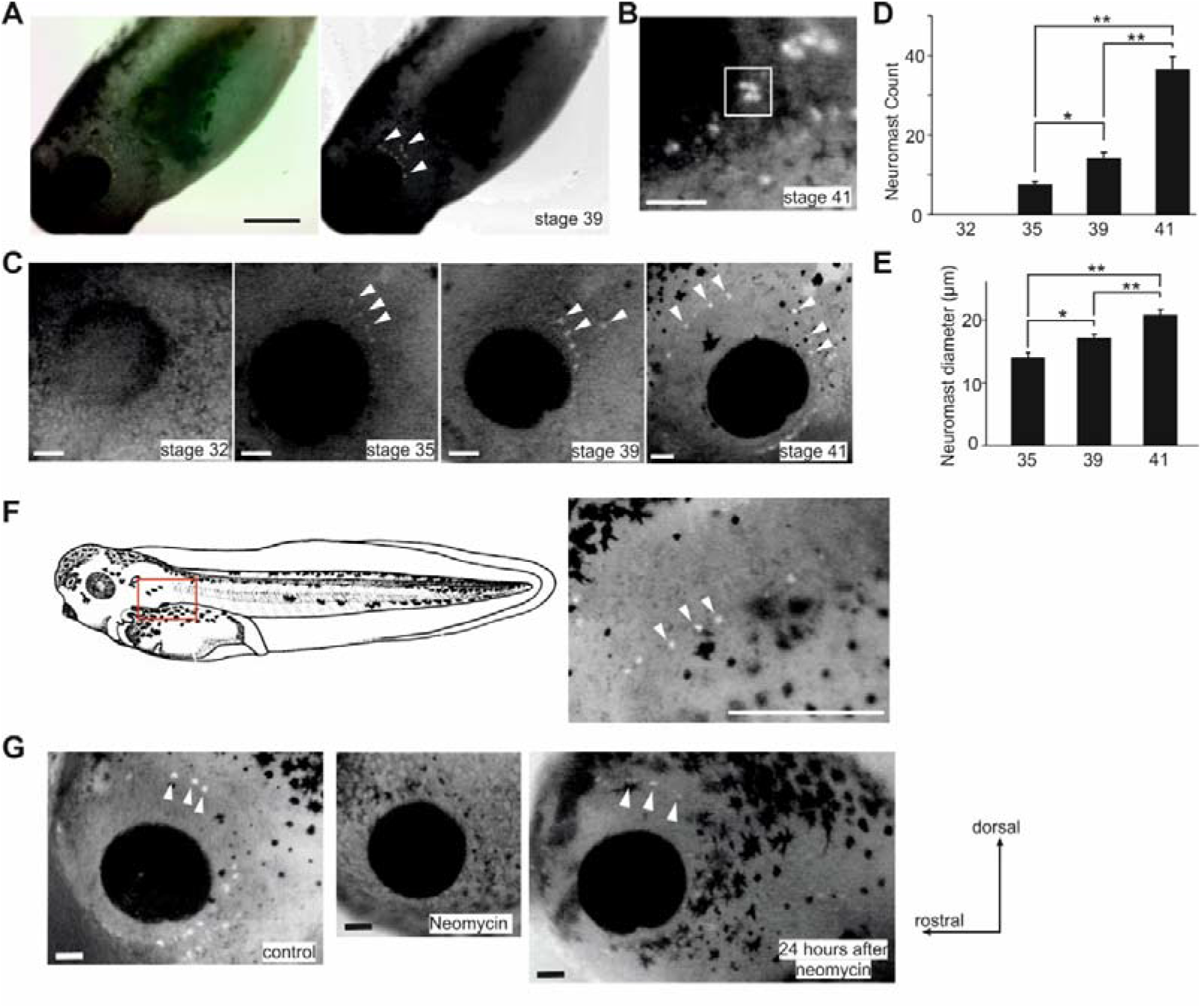
Staining tadpole lateral line neuromasts using DASPEI in control and after neomycin treatment. **A.** Converting a colour photo of a tadpole after DASPEI staining to grey scale after enhancing the red and yellow channels. **B.** Clustering of neuromasts in a stage 41 tadpole (square). **C**. DASPEI staining of tadpoles at various stages (only eye region shown). D. Neuromast counts for each side of tadpoles at stages 32, 35, 39 and 41. **E.** Neuromast diameter in stage 35, 39 and 41 tadpoles. **F.** The appearance of posterior LL neuromasts. Photo shows the red rectangle area in the drawing on the left. **G.** DASPEI staining of stage 40 tadpoles in control, treatment with 3 μM neomycin for 30 minutes and 24 hours after neomycin treatment. Orientation of the tadpoles is the same in **B, C** and **G.** White arrow heads point at some example neuromasts labelled with DASPEI. Scale bars: 500 μm in **A** and **F**; 100 μm in **B, C** and **G**.

Neomycin was obtained from Sigma-Aldrich UK in solution (10 mg/ml in 0.9% NaCl solution) and stored at 4°C. To destroy the neuromast hair cells, tadpoles were placed in a 3 μg/ml solution of neomycin in saline for 30 minutes. Tadpoles were then imaged to confirm neuromast ablation (Roberts A, Feetham B, Pajak M and Teare T, 2009).

Before testing tadpole responses to suction, the animal tail was touched with a hair to ensure it was capable of swimming normally. A modified water pump was used to apply suction through plastic tubing (inner diameter: ~ 1.5 mm), with the end of the tube placed ~ 1 mm from the head of the tadpole. The escape response of each tadpole was tested five times, with a rest period of three minutes between trials. For motor nerve (m.n.) recordings, tadpoles were immobilised by α-bungarotoxin at 10 μM for 20-30 minutes and dissections were made to expose the muscle cleft. A smaller nozzle with a diameter of ~ 120 μm was placed ~ 150-200 μm above the left eyecup. The suction level was set by manually changing the volume of air trapped in a stock bottle of 500 ml using a 50ml syringe. Suction level was calculated based on Boyle’s law (P1xV1=P2xV2) and expressed as the difference from normal atmosphere pressure in KPa. The application of suction for the duration of 0. 5-10 seconds was controlled by a Toohey spritzer (Toohey Company, Fairfield, NJ, USA), the valve of which connects/disconnects the bottle with the suction nozzle. Extracellular recordings of motor nerve activity were made using methods described previously (Li WC et al., 2017). Briefly anaesthetised and then immobilised tadpoles were dissected to remove the yolk belly and expose the swimming myotomes in the trunk region. Glass electrodes filled with saline were placed on one or two muscle clefts (normally between the 5^th^ and 6^th^ myotomes). Recording of anterior LL nerve (aLLN) activity was carried out by positioning one glass electrode on the cut nerve end with gentle suction (~-200 Pa). LED dimming, electrical skin stimulation and suction application were all controlled by TTL pulses configured in the sampling software Signal, through the power 1401 board (CED, Cambridge, UK). The suction was monitored using a fluid flow sensor to monitor if the suction nozzle was clogged during experiments.

In order to image the intracellular calcium signals in brainstem neurons, some ependymal cells inside the brainstem were removed to expose neuronal somata by dissection (Li WC, Zhu XY and Ritson E, 2017). Tadpoles were left in 5 μM Fluo-4 AM saline solution for ~20 minutes in darkness. After resting the tadpole for about 20 minutes, fluorescence images were captured at 5 Hz using x10 or x20 water immersion lenses with a Neo5.5 CMOS camera and the Andor Solis software (Oxford Instruments, UK).

Some processing of the m.n. recordings was conducted in Dataview, courtesy of Dr William Heitler at the University of St Andrews. All data sets were examined for normality first. Non-parametric statistical methods were used for those without normal distributions using IBM SPSS.

## Results

### DASPEI staining in control and after neomycin treatment

We first aimed to count and locate the LL neuromasts using fluorescent DASPEI staining in live tadpoles at different developmental stages. No neuromast was visible at stage 32. Most neuromasts appeared as lone spots at stage 35 and 39 with some appearing to cluster together especially at stage 41. In the latter case, the number of separate dots in the clusters was often difficult to resolve and they were counted as one neuromast (Fig. 1B). The mean neuromast count increased progressively, rising from 7.6 ± 2.46 per tadpole at stage 35 to 14.2 ± 4.39 at stage 39, and 36.5 ± 10.02 at stage 41 (n = 10 tadpoles for each stage, *p* < 0.001, one way ANOVA, Fig.1C-D). The mean diameter of stained neuromasts also increased significantly over the stages examined (Figure 1E), from 13.92 ± 3.15 μm at stage 35, to 17.12 ± 2.71 μm at stage 39, and finally to 20.72 ± 3.9 μm at stage 41 (n = 15 neuromasts for each stage, *p* < 0.001, one way ANOVA, Fig. 1E). Until stage 39, neuromasts were only observed on the head, mostly arranged in a single line caudal to the eye. At stage 39, in some cases new neuromasts began to appear posterior to the original neuromast line (Figure 1A). By stage 41, neuromasts were observed surrounding the eye, extending to the dorsal area (Figure 1C). At this stage the posterior LL neuromasts also started to appear in a single line down the trunk, but never reaching the tail (Fig. 1 F).

We next tested tadpole responses to suction at rest at development stage 40. Stage 40 tadpoles had, on average, a 61.8 ± 4.2% probability of escaping the suction applied (n = 11 tadpoles, 5 trials each), i. e. they produced a C-bend of their trunk and then swam away from the suction nozzle, or turned inside the suction nozzle and swam out. In order to verify if this escape response was initiated by the activation of tadpole LL system, the aminoglycoside antibiotic, neomycin, was used to destroy the neuromasts. After exposing tadpoles to 3 mg/ml neomycin for 30 minutes, DASPEI staining did not label any neuromasts (n = 5 tadpoles, Fig. 1G). Neomycin-exposed tadpoles also lost their escape response to suction, i.e. they were passively sucked in the nozzle without motor responses like C-bend and swimming. Their escape probability decreased from 60 ± 6.3% to 0% and showed no recovery 24 hours after transferring to normal saline (5 trials to each tadpole in every condition) although tadpoles responded to touch stimulation with swimming reliably (100%, 5 trials in each condition) immediately after neomycin treatment and after 24-hour rest period. After the 24-hour rest period, a mean of 28 ± 5.25 neuromasts from both left and right sides were detected using DASPEI labelling for the second time (n = 5 tadpoles), indicating some regeneration of the LL neuromasts. The location of these neuromasts was similar to that in control tadpoles but staining did not appear as intense (Fig. 1G).

### Activating the lateral line system in immobilised tadpoles

Having located the neuromasts, a suction nozzle could be positioned in their vicinity to activate them and investigate the effect on tadpole motor outputs. We recorded tadpole motor outputs in immobilised tadpoles to determine whether we could replicate the escape response (Fig.2). The tadpole was pinned on its side onto the sylgard block in the recording dish, similar to the position tadpoles assume when they rest at the bottom of a petri dish. A nozzle with a diameter of ~120 μm was placed within ~150-200 μm above the left eyecup and used to apply suction (0.5-10 seconds, −1 to −7.5KPa).

**Fig. 2.**
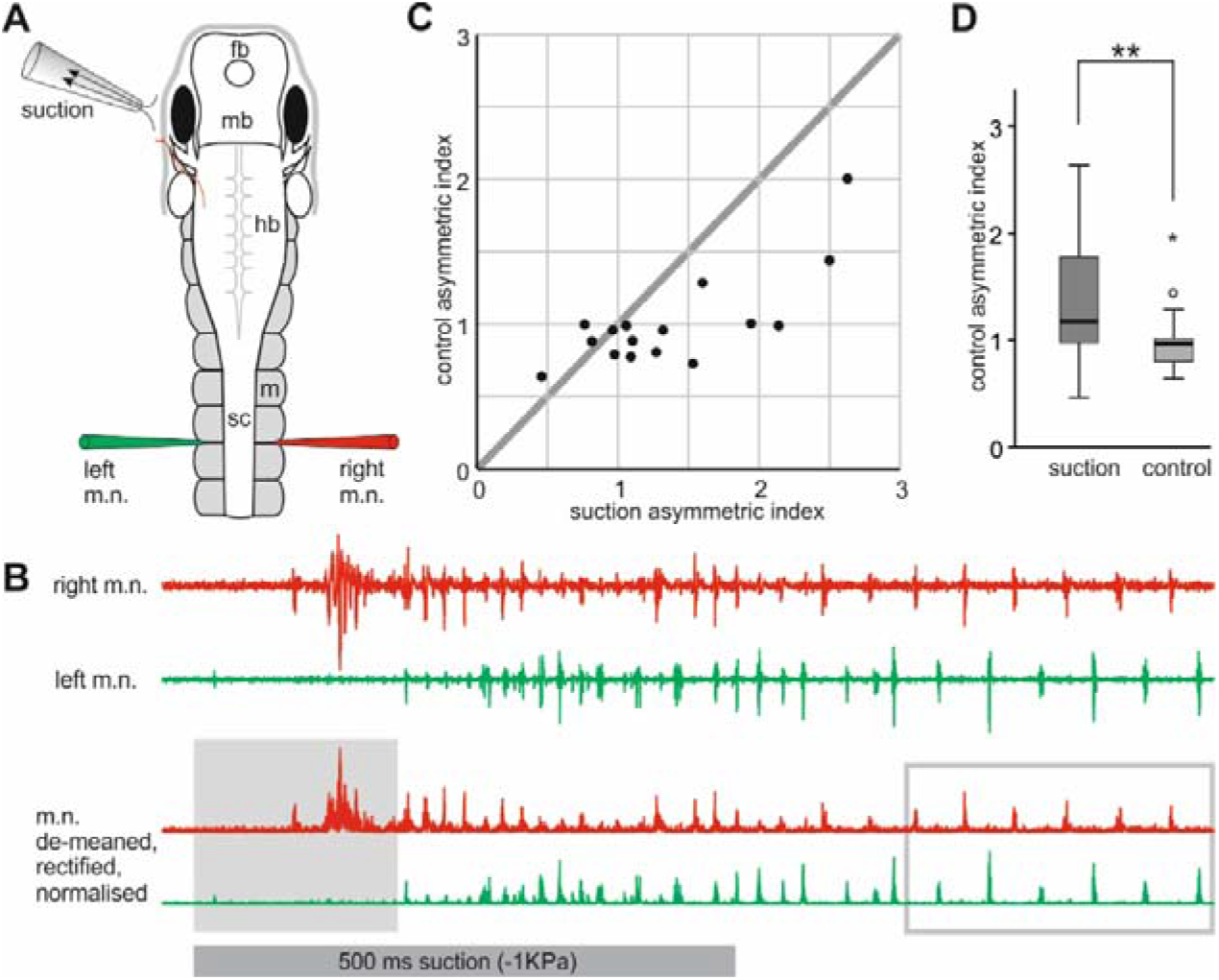
Asymmetric motor responses elicited by local suction close to the left eyecup. **A**. Experimental setup diagram showing tadpole anatomy and the position of the suction pipette and recording electrodes. fb, forebrain; mb, midbrain; hb, hindbrain; sc, spinal cord; m, myotome. **B.** One example initial motor response elicited by a 500ms suction pulse (grey bar below) and its processing for calculating asymmetric index. Grey area includes the period of de-meaned, rectified and normalised recordings for integration. Grey rectangle encircles a few swimming cycles where m.n. bursts are used for normalising m.n. activity. Traces are color-coded to electrodes. There is a lot of synchrony (Li WC, Merrison-Hort R, Zhang HY and Borisyuk R, 2014) before swimming. **C.** The average asymmetric indices in control, spontaneous swimming is correlated with those in motor responses induced by suction. Thick grey line is the identity line. **D.** Asymmetric indices are larger in suction-evoked responses than in control. ** shows significance at *p* < 0.01.

Suction often elicited asymmetric motor responses between the two sides monitored with simultaneous left and right motor nerve recordings (Fig.2A). Using Dataview software, we first reset the baseline m.n. recording at 0 (de-meaning), rectified the traces and integrated the m.n. activity for the period starting from the onset of suction to the point when rhythmic m.n. bursts appeared (either synchrony or swimming, Fig.2B, (Li WC et al., 2014)). Assuming the activity between the two sides will be symmetrical during swimming, we normalised the amplitude of m.n. activity to the bursts during the first 15-20 swimming cycles. We then calculated the asymmetric index by dividing the normalised and integrated m.n. activity on the right side by that on the stimulated, left side. This was calculated for 5 suction-evoked responses and 5-8 spontaneous swimming episodes in each tadpole and averaged. There was a correlation between the control average asymmetric indices and those for suction (Spearman’s rank correlation, *p* < 0.05, Fig.2C). The correlation coefficient of 0.562 suggests that animals that showed tendency to bend away from the suction side at the beginning of spontaneous swimming produced clearer escape response to suction. The average indices were higher in the case of suction (1.39 ± 0.16) than in control (1.01 ± 0.08, *n* = 16 tadpoles, related sample Wilcoxon Signed rank test, *p* < 0.01, Fig.2D), suggesting the tadpole would initially bend its body away from the suction, followed by swimming.

We observed that when suction was applied in the middle of on-going swimming activity swimming was halted (Fig.3A, B). Normal swimming episodes were 10 – 41 seconds long (n =12 tadpoles). We applied a 0.5 second suction when the swimming was about halfway through its episode in control conditions (8-12 trials in each tadpole). The suction reliably shortened swimming in 127 out of 131 trials (independent sample t-test, all *p* < 0.05, Fig.3C) which lasted for 1.32 ± 0.16 seconds after suction ended.

**Fig. 3.**
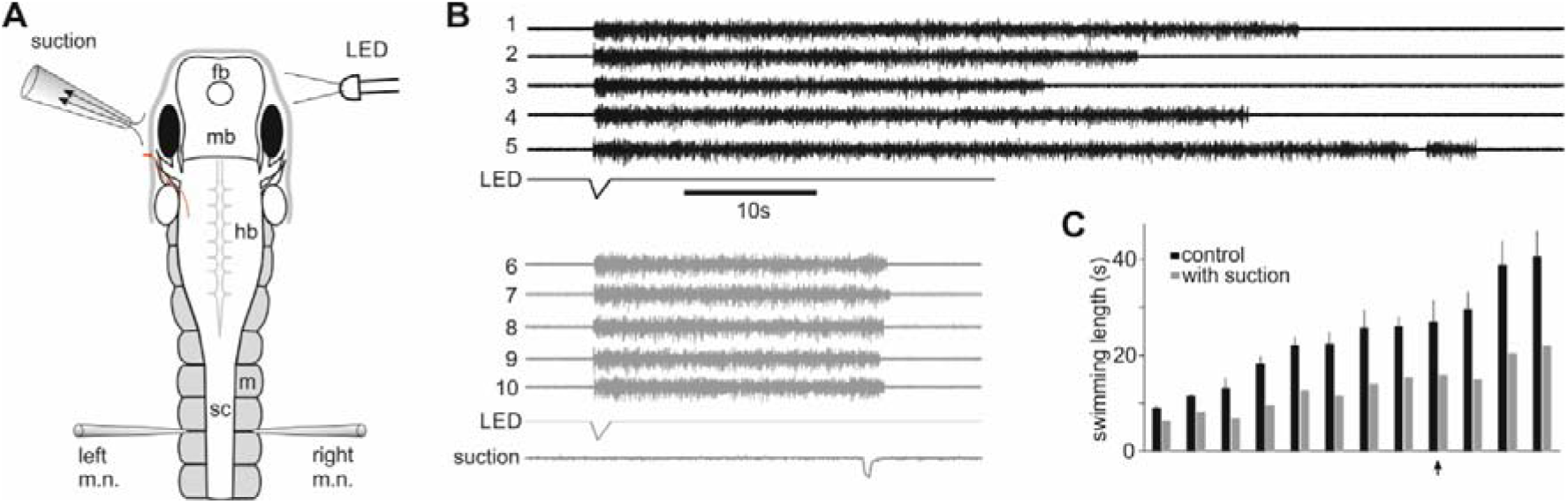
Suction stops on-going swimming. **A.** Experimental setup showing tadpole anatomy and the position of the LED light, suction pipette and recording electrodes. For abbreviations see Fig.1A. **B**. Consecutive swimming episodes initiated by dimming the LED light in control (trials 1-5) and with 500ms suction applied ~ 20 seconds into swimming (trials 6-10). Only activity from the left m.n. is shown. **C.** Summary of swimming lengths after suction in 12 individual tadpoles (all *p* < 0.01 except *p* = 0.012 in the tadpole with an arrow).

### The effects of electrical stimulation of the anterior lateral line nerve

Activating the lateral system using suction was most effective when minimal dissection was made to the tadpoles. In order to study the neuronal pathways in the central nervous system that mediated these motor responses, we needed to open up the brainstem. We next tried to stimulate the anterior LL nerve (aLLN) to see if we could reproduce the motor response above. We cut the aLLN connecting to the head skin using a pair of fine scissors and used a stimulation electrode (diameter: ~ 60 μm) to suck onto the severed end of aLLN. One to 20 stimulation pulses (0.5 −2ms in duration, 200-400 Hz) were used to excite the aLLN nerve. Following stimulation, swimming was initiated reliably but without clear prolonged one-sided bursts indicating an escape response (Fig.4B). Therefore, we did not attempt to calculate the asymmetric indices. When stimulation was applied in the middle of on-going swimming, swimming could be stopped reliably (Fig.C, D), just as with the more natural local suction stimuli. Swimming after stimulation lasted for 3.5 ± 0.3 s, much shorter than in control conditions (47.5 ± 5.4s, *p* < 0.001, paired t-test, 68 trials in 9 tadpoles).

**Fig. 4.**
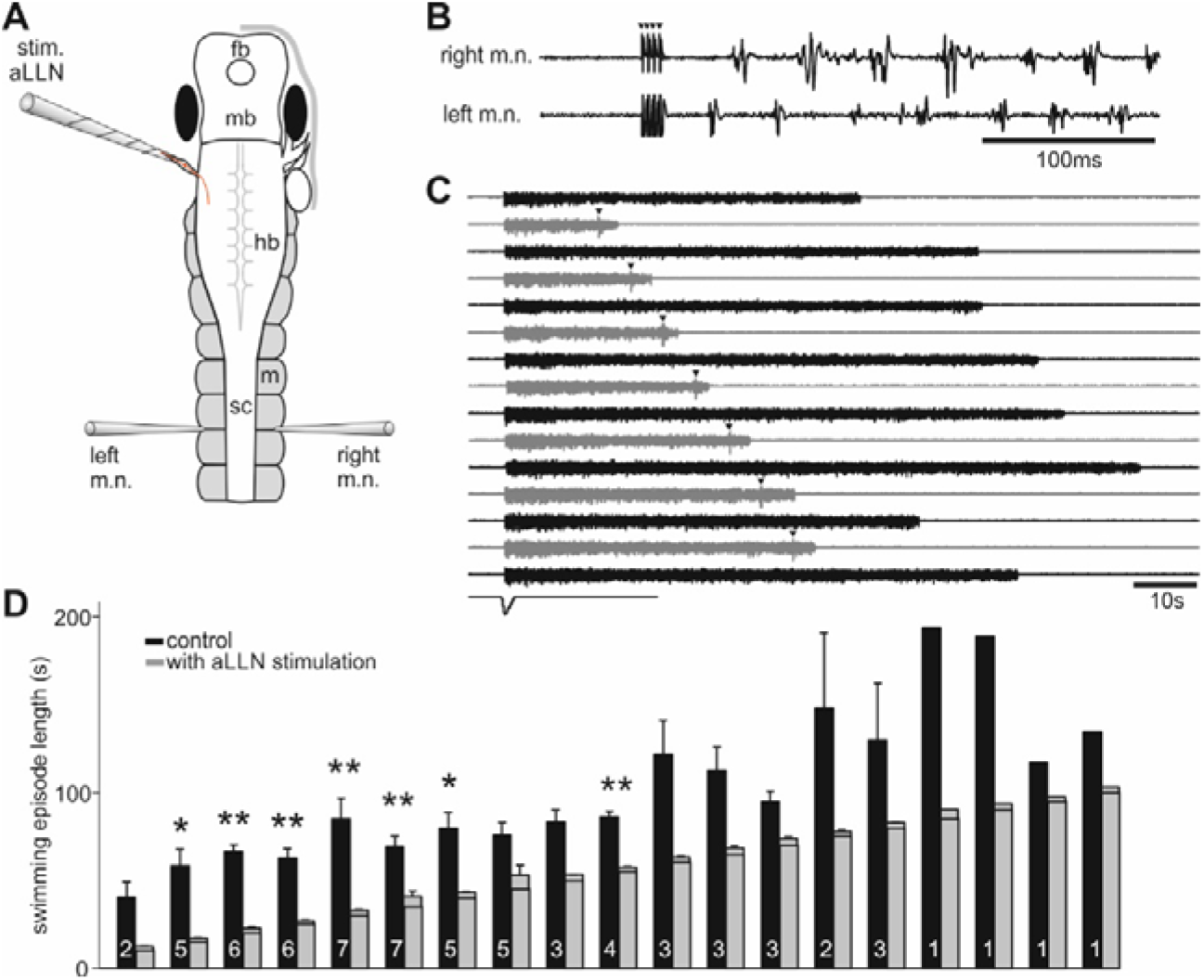
Stimulating the anterior lateral line nerve (stim. aLLN) electrically does not produce clear escape response but stops on-going swimming. **A.** Experimental setup showing tadpole anatomy and the position of the stimulation and recording electrodes. **B.** An example of stimulating aLLN repetitively (4 x 0.5 ms pulses at 20 μA and 200 Hz, arrow heads) not evoking prolonged escape burst before alternating swimming bursts between the left and right m.n. recordings. **C.** Single electrical stimulation (arrow heads) applied at various points after swimming initiation stops swimming reliably (grey traces, black is control). **D.** Summary of electrical stimulation of aLLN at different time from the beginning of swimming (lines within the grey bars) shortening swimming (* indicates significance at p<0.05, ** at p<0.01, paired t-tests). Numerals inside black control bars are number of tadpoles tested for each time point of electrical stimulation.

### Afferent and efferent aLLN activity

We next recorded the afferent and efferent activities of the aLLN. aLLN was severed at its entry point to the hindbrain. A suction pipette with ~ 60 μm tip diameter sucked onto the cut end while maintaining the skin around the eye intact. A suction nozzle was placed close to the eyecup to activate the aLLN peripherals (typical duration: 7 seconds, Fig.5A, B). In most cases, multiple units were recorded often with similar amplitudes in their extracellular action potentials. To simplify analyses, we did not try to discriminate different units. Instead, we used these extracellular discharges to trigger events once they reached the threshold (arbitrarily set at ± 5 SD of the baseline, Fig.5B). Then we counted the number of events in each 0.5 second bin and averaged them across 14 tadpoles. The average number of events was higher during the suction period than before or after suction (p < 0.01, related-samples Wilcoxon signed rank test). The latency from the start of suction flow to the first unitary discharge was 19.7 ± 1.1 ms (13 trials in 13 tadpoles). The discharges to suction also showed typical adaptation with time (Fig.5C).

**Fig. 5.**
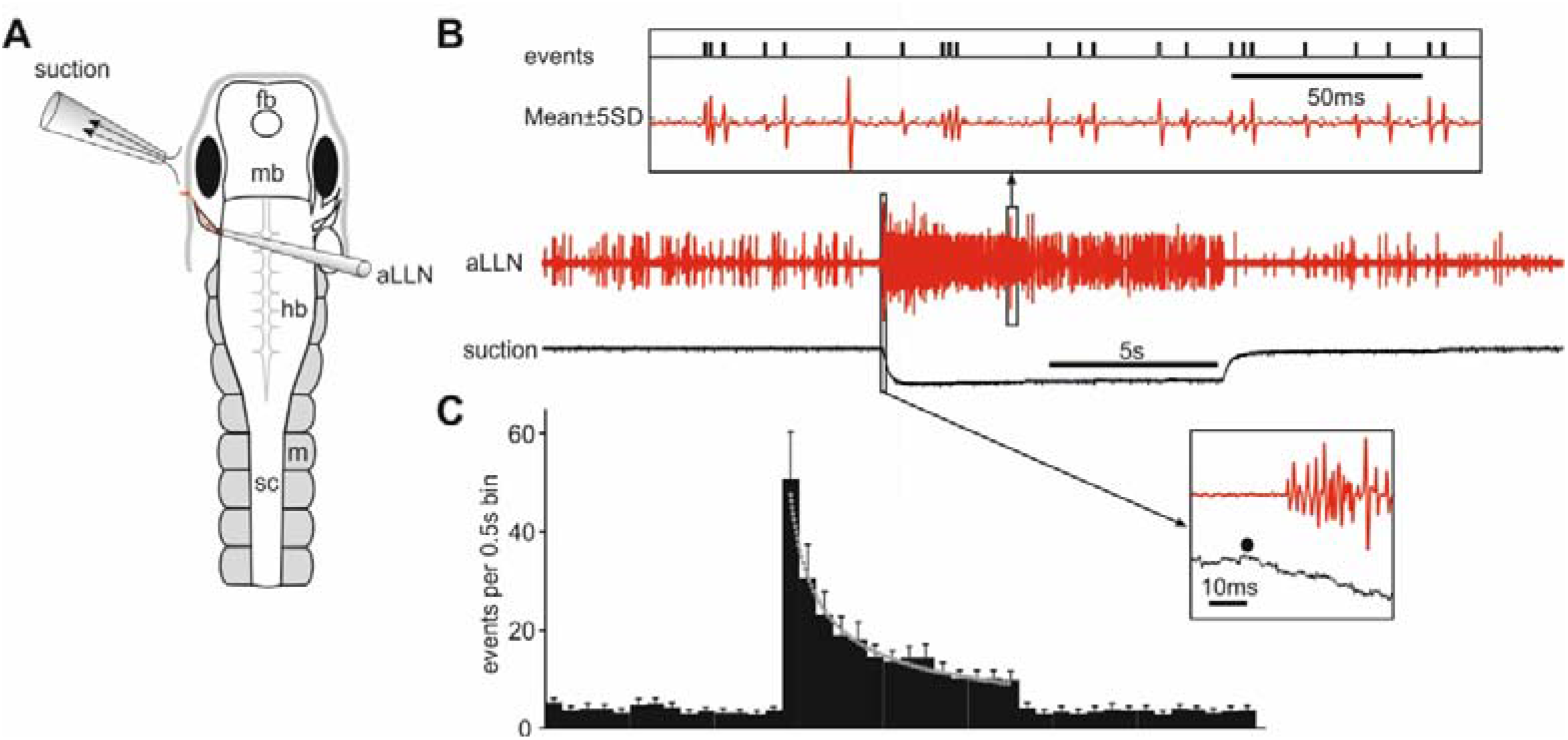
aLLN afferent activities. **A.** Experimental setup for recording aLLN afferent activity. The recording electrode directly sucks onto the cut end of aLLN. **B.** An example recording of aLLN activity around time of 10s suction (−3.3 KPa). Inset above (boxed area) shows events triggering by setting the threshold at ± 5SD in the aLLN recording trace. Inset below shows the delay from the beginning of saline flow (dot) to first unitary discharge. **C.** Average binned events in 14 tadpoles showing increased activity during a 7s suction period and adaptation (bin width: 0.5s). Dotted fitting curve is for activity during suction: y = 47.33 X^-0.61^ (R^2^ = 0.98).

We removed the skin covering the left eyecup and cut the distal end of aLLN using a pair of fine scissors. The otic capsule was also exposed and removed, together with the trigeminal nerve. A pipette with a diameter of ~ 60 μm was used to suck onto the cut end to record aLLN efferent activity with m.n. activity recorded simultaneously (Fig.6A). Efferent activity was only recorded during swimming, which was initiated by dimming an LED light (Fig.6B). The efferent discharges were more reliable at the beginning of swimming episodes and became unreliable a few seconds after swimming was started (Fig.6C). The distribution of efferent discharge timing showed an in-phase peak with ipsilateral (left) m.n. bursts (Fig.6D).

**Fig. 6.**
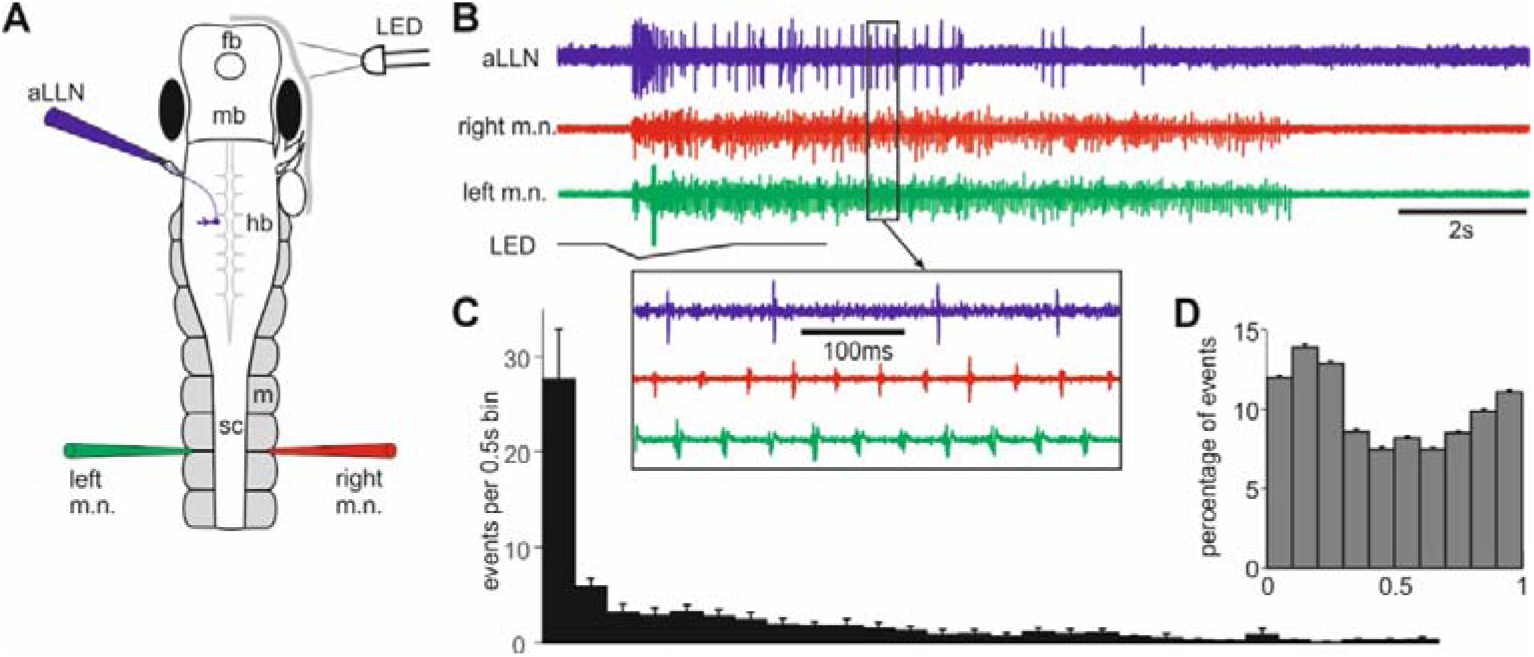
aLLN efferent activities. **A. Setup for recording aLLN efferent activity where swimming is started by dimming an LED light. B.** An example aLLN efferent recording during a swimming episode. Box region is expanded to show timing of the efferent activity relative to m.n. bursts on both sides. **C.** Number of unitary discharges per 0.5s bin in the first 14 seconds after swimming is started (averaged from 3 episodes from each of 9 tadpoles). **D.** Phase of aLLN unitary discharges plotted against the m.n. bursts on the ipsilateral side (100 spikes from each of 10 tadpoles).

### Locating lateral line sensory interneurons using calcium imaging

In order to trace the LL pathways in the central nervous system, we used calcium imaging to locate the sensory interneurons in the hindbrain on the stimulated side. We opened the dorsal roof of the brain stem and removed some ependymal cells lining the inside wall of hindbrain and midbrain so Fluo-4 AM could be loaded into the exposed neurons. After this dissection, the stub of any cut aLLN would be obscured by the hindbrain which opened up sideways when viewed from the top. This made it impractical to electrically stimulate the aLLN. Instead, we positioned a suction nozzle with a tip diameter of ~120 μm close to the left eyecup to activate aLLN. A 10x water immersion objective was used so a large area of hindbrain and midbrain could be imaged to screen active neurons. Simultaneous m.n. nerve and suction flow recordings were carried out (Fig.7).

**Fig. 7.**
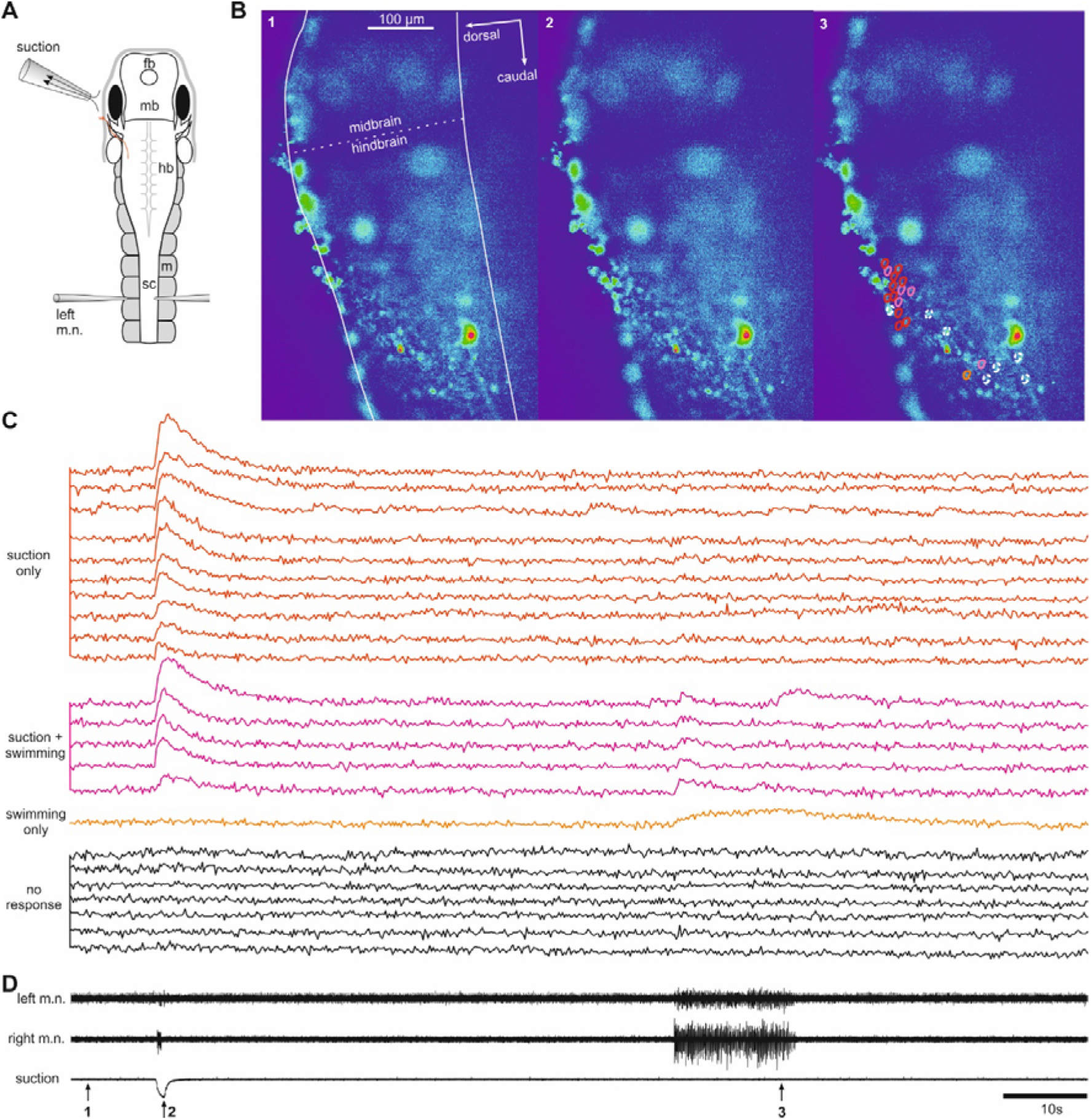
Locating aLLN sensory interneurons using calcium imaging. **A.** Experimental setup for calcium imaging using a x10 water immersion objective. The preparation is tilted so the left side of the hindbrain stays roughly flat to facilitate imaging many neurons in a single focal plane. Imaging is at 5 Hz for 120s. **B.** Three frames captured at the indicated time points in **D** (1,2 and 3). Lines in frame 1 shows the profile of the hindbrain and midbrain (border: dashed line). Circles in frame 3 indicate the location of neurons whose calcium activities are given in **C. C.** Calcium activity of 24 neurons outlined by circles in **B** (color-coded except that dashed white circles correspond to black traces). Different types of responses are grouped as labelled. **D.** Simultaneous m.n. and suction recordings.

In 6 tadpoles, we located neurons with increased calcium activities immediately after suction (0.5 s duration) in the very dorsal hindbrain region where the aLLN entered the hindbrain (49 Neurons, e.g. see red and pink traces in Fig.7C). In two of the six tadpoles, suction did not evoke any motor response but still activated 6 and 5 neurons in this region. In three of the six tadpoles where spontaneous swimming was also monitored, 13 out of the 35 aLLN neurons were also active at the start of spontaneous swimming. However, such activity did not last through the swimming episode (pink traces in Fig.7C). These data suggest that the identified 49 neurons are sensory interneurons for the aLLN.

## Discussion

In this study, we have examined the basic physiology of aLLN in immobilised tadpoles after locating the neuromasts using DASPEI labelling. The results showed that the aLLN plays a role in potential tadpole escape behaviour, initiation of swimming and termination of on-going swimming.

In adult amphibians, each LL stitch is innervated by two myelinated afferent fibres (Kroese AB, Van der Zalm JM and Van den Bercken J, 1978; Nagiel A, Andor-Ardo D and Hudspeth AJ, 2008; Strelioff D and Honrubia V, 1978), projecting along the lateral spinal tract to a specialized hindbrain nucleus (Roberts A, Feetham B, Pajak M and Teare T, 2009; Schlosser G, 2002). Each afferent is excited by one orientation of hair-cell, and inhibited by the other (Kroese AB, Van der Zalm JM and Van den Bercken J, 1978; Obholzer N et al., 2008; Strelioff D and Honrubia V, 1978). Afferent neurons of the head receptors project to the aLLN, and trunk receptors to the posterior LL nerve (Northcutt RG, 1992).

What are the primary functions for the LL system in young *Xenopus* tadpoles? Rheotaxis is particularly important in river-dwelling species and those inhabiting turbulent water to prevent being swept away. CoCl_2_ ablates superficial neuromasts, significantly increasing the current threshold for rheotaxis in fish (Montgomery JC, Baker CF and Carton AG, 1997), and impairing the ability of older *Xenopus* tadpoles to orient to a current source (Simmons AM, Costa LM and Gerstein HB, 2004). In tadpoles, directional orientation (facing into the current) is observed at stages 37-46 (Nieuwkoop PD and Faber J, 1956; Simmons AM, Costa LM and Gerstein HB, 2004), but positive rheotaxis (actively swimming against the current) is not observed until stage 47 (Nieuwkoop PD and Faber J, 1956; Roberts A, Feetham B, Pajak M and Teare T, 2009). The hatchling *Xenopus* tadpole at stage 37/38 is inactive 99% of the time, in comparison to constant swimming at later stages. This suggest that rheotaxis may not be a primary function of the LL system. The initial asymmetrical motor nerve bursts recorded in immobilised tadpoles could signal a turning in body orientation. This turning behaviour is necessary for aquatic prey animals to swim against water flow generated by inertial suction-feeding predators (Carreno CA and Nishikawa KC, 2010). In zebrafish, this is typically the C-start escape response, allowing larvae to escape 68% of simulated strikes (McHenry MJ, Feitl KE, Strother JA and Van Trump WJ, 2009) and 70% of attempted predatory strikes by the adult zebrafish (Stewart WJ, Cardenas GS and McHenry MJ, 2013). Similar behaviour has been seen in tadpoles (Roberts A, Feetham B, Pajak M and Teare T, 2009). Ablating the hair cells with neomycin prevented escape responses in zebrafish (McHenry MJ, Feitl KE, Strother JA and Van Trump WJ, 2009; Stewart WJ, Cardenas GS and McHenry MJ, 2013) and tadpoles as shown in this study. The C-start in fish is mediated by Mauthner neurons, but the directionality of this response is dependent on aLLN input (Mirjany M and Faber DS, 2011). Mauthner functionality in tadpoles is uncertain, but there is suggestion for their activity (and a full C-start response) at stage 42 (Sillar KT and Robertson RM, 2009). The turning in young tadpoles should involve neuronal pathways other than Mauthner neurons, as *Xenopus* tadpoles can react to water current as early as stage 31/32, coinciding with the appearance of the first neuromasts (Roberts A, Feetham B, Pajak M and Teare T, 2009). LL function in post-metamorphosis *Xenopus* changes from predator evasion to detection of prey (Claas B and Munz H, 1996), when it predates using the same inertial suction feeding mechanism it must avoid in its larval stage (Carreno CA and Nishikawa KC, 2010). Opposing hair-cell populations can detect water flow in two axes: anterior-posterior, or more rarely, dorsal-ventral (Nagiel A, Andor-Ardo D and Hudspeth AJ, 2008), allowing coding information in four directions. Precise directional information can be relayed when all afferent activities from different neuromasts in the animal body are processed in the CNS. It is likely that the use of a narrow suction nozzle in our experiments reduced the chance of a full reproduction of escape response in immobilised tadpoles as in control, although asymmetrical motor discharges were seen in most tadpoles.

Apart from the role in escape responses, we have revealed that activating the LL system could unexpectedly stop on-going swimming. This opposing effect on motor output depending on whether the tadpole is active or not is similar to the response reversal observed in other preparations (Chase MH and Wills N, 1979; Pearson KG and Collins DF, 1993). Such reversed response was also seen when the head skin was stimulated, which could evoke swimming at rest but terminate on-going swimming (Li WC, Zhu XY and Ritson E, 2017). It will be interesting to see if the LL system and the touch sense in the head skin share the same neuronal pathways in terminating swimming.

There has been significant progress on how mechanical stimuli are transduced to electrical signals in the LL afferents, especially in the posterior LL system in fish. Successful whole-cell patch-clamping has allowed the ion-channel activity and glutamate release from hair-cells to be observed *in vivo* (Ricci AJ, Bai JP, Song L, Lv C, Zenisek D and Santos-Sacchi J, 2013). Mechanoelectrical transducer cation channels (Fettiplace R, 2009) on the hair-cells open or close with kinocilium deflection caused by water flow and, together with the inherent outward K+ current (Bleckmann H and Zelick R, 2009; Fettiplace R, 2009; Ricci AJ, Bai JP, Song L, Lv C, Zenisek D and Santos-Sacchi J, 2013), control glutamate release from the hair-cell base (Fettiplace R, 2009; Ricci AJ, Bai JP, Song L, Lv C, Zenisek D and Santos-Sacchi J, 2013; Trapani JG and Nicolson T, 2011). The role of glutamate is supported by the necessity of vesicular glutamate transporter (VGluT3), which is exclusive to the basal region of inner ear and LL hair-cells. Zebrafish *Asteroid* mutants without VGluT exhibit no afferent LL action potentials (Obholzer N, Wolfson S, Trapani JG, Mo W, Nechiporuk A, Busch-Nentwich E, Seiler C, Sidi S, Sollner C, Duncan RN, Boehland A and Nicolson T, 2008). Afferent neurons of the LL exhibit irregular spontaneous spiking (Kroese AB, Van der Zalm JM and Van den Bercken J, 1978; Trapani JG and Nicolson T, 2011), which may result from voltage-activated calcium channel-mediated baseline release of glutamate from the hair-cells (Trapani JG and Nicolson T, 2011). Our study here also revealed spontaneous afferent activities in the tadpole aLLN. Further studies are needed to see if similar ionic mechanisms sustain these spontaneous activities.

In terms of LL efferents, it has been shown that both cholinergic and dopaminergic efferent neurons modulate the hair-cell output (Haehnel-Taguchi M, Akanyeti O and Liao JC, 2018). In stages 48-55 tadpoles, the efferent activity was shown to be corollary to the locomotor rhythms, in phase with ipsilateral spinal central pattern generator activity and suppressing afferent sensory signalling (Chagnaud BP et al., 2015). These efferent neurons are located in the brainstem nucleus which projects to innervate both lateral line and inner ear hair cells (Hellmann B and Fritzsch B, 1996).

There are some unidentified cholinergic neurons in tadpole brainstem, which are involved in termination of swimming in the concussion-like events (Li WC, Zhu XY and Ritson E, 2017). We do not know if these cholinergic neurons are also candidates of the cholinergic efferents in the tadpole LL nerves. Whilst cholinergic modulation is inhibitory (Bricaud O et al., 2001), the role of dopaminergic modulation is not understood. In stage 37/38 tadpoles, the monoaminergic systems including dopaminergic modulation are still not endogenously functional (Sillar KT et al., 2014). The efferent activity therefore unlikely represent dopaminergic modulation.

In fishes, afferent LL activity propagates from the peripheral end of afferents and enters the hindbrain to the caudal and medial octavolateralis nucleus (Maruska KP and Tricas TC, 2009). In adult *Xenopus laevis*, the aLLN projects to a longitudinally dispersed area in the brainstem which includes the LL nucleus and the medial part of the anterior nucleus (Will U et al., 1985). Similar to that in hagfish, *Xenopus* aLLN ganglion is closely attached to the trigeminal ganglion (Kishida R et al., 1987). It is not clear if the aLLN consists purely of LL sensory cells or a mixture of cutaneous sensory cells and lateral line sensory cells as in the hagfish, where projections run to both the trigeminal sensory nucleus and medial nucleus of the area acousticolateralis. Using calcium imaging, we have located a number of sensory interneurons responding to suction stimulation with short latencies. Most of these neurons are packed within the region likely corresponding to the lateral line nucleus. Even within this region, some neurons appeared to be also involved in the initiation of swimming activity whilst others only showed activity in response to suction. Future studies need to trace where these neurons project to in the central nervous system and how the LL sensory information is further processed to lead to various motor outputs including escape responses, initiation and termination of swimming and rheotaxis.

## Acknowledgement

Valentina Saccomanno was an exchange MSc student in the University of St Andrews with a scholarship by the Erasmus+ Programme for Traineeship and the University of Trieste. Heather Love and Amy Sylvester were undergraduate neuroscience students conducting their senior honours research projects in the University of St Andrews.

